# VirTrack: A Framework for Inferring Viral Influence on Disease-Associated Transcriptomes — Clinical Type-Specific Epstein–Barr Virus Pathogenesis in Multiple Sclerosis

**DOI:** 10.1101/2025.10.05.680499

**Authors:** Anna Onisiforou

**Author notes:** <u>**Corresponding author:**</u> Anna Onisiforou, Ph.D., Senior Scientist and Head of the AI & Systems Bioinformatics Unit, Department of Psychology, University of Cyprus, 75 Kallipoleos Avenue, Aglantzia, Nicosia 1678.

## Abstract

**Background:** Epstein–Barr virus (EBV) is strongly implicated in Multiple Sclerosis (MS), but how its influence varies across MS clinical types remains unclear.

**Methods:** We developed *VirTrack*, a computational framework that integrates experimentally validated EBV–host protein–protein interactions (PPIs) with clinical type–specific peripheral blood transcriptomes from Clinically Isolated Syndrome (CIS), Relapsing Remitting MS (RRMS), Secondary Progressive MS (SPMS), and Primary Progressive MS (PPMS) (GSE136411). *VirTrack* (i) anchors viral interactions in differentially expressed genes (DEGs), (ii) ranks EBV targeting of DEGs and host hubs, and (iii) applies machine learning– based clustering to identify clinical type–specific functional pathways influenced by EBV.

**Results:** EBV engagement was clinical type–dependent. In early MS (CIS and RRMS), EBV targeted approximately 13–18% of dysregulated genes, enriching for B-cell–related processes, Toll-like receptor signaling, and infection-like inflammatory pathways, while suppressing antiviral and NF-κB responses. Progressive clinical types exhibited fewer viral connections but distinct mechanistic shifts: SPMS was characterized by suppression of vascular and cardiac repair–associated pathways, whereas PPMS was dominated by upregulation of vacuolar and lysosomal remodeling processes. Hub analyses revealed a stable core of influential EBV proteins, EBNA-LP (consistently top-ranked), BZLF1, BVLF1, LMP2, and BDLF4, while BNLF2A showed broad but less hub-focused targeting and LMP1 ranked low across types.

**Conclusion:** EBV shapes MS through dynamic, clinical type–specific perturbations, driving strong immunomodulation in early disease and selective cellular remodeling during progressive types. *VirTrack* identifies key viral proteins and host pathways for stage-tailored therapeutic targeting, supporting early EBV-directed interventions and revealing potential links to vascular comorbidity in progressive MS types. More broadly, *VirTrack* offers a generalizable, systems-level framework for elucidating viral contributions across complex human diseases.

## 1. Introduction

Viruses are increasingly recognized as key environmental factors that contribute to the onset and progression of complex human diseases, including neurodegenerative disorders, autoimmune diseases, and cancers ^1–5^. They can act as triggers of disease initiation and/or progression, amplifiers of immune dysregulation, and modulators of host molecular networks. Among these, Epstein–Barr virus (EBV), a ubiquitous herpesvirus infecting more than 90% of the human population, has emerged as a leading candidate in the pathogenesis of Multiple Sclerosis (MS) ^6,7^.

A recent pivotal longitudinal study demonstrated that individuals who seroconverted to EBV had more than a 30-fold higher likelihood of developing MS, and almost all MS patients show evidence of prior EBV infection, supporting the causal role of EBV in MS ^6^. Furthermore, elevated EBV-specific antibody titers and altered immune responses against latent viral proteins have been observed in MS patients ^8^. While recent evidence strongly supports the central role of EBV as a trigger for MS, debate remains as to whether EBV is universally causal or one of several environmental factors contributing to disease initiation. Even if EBV is not the sole initiating agent, its persistent presence may act as a cofactor that amplifies autoimmune responses and sustains disease progression. Therefore, regardless of its role in initiation, targeting EBV remains a relevant therapeutic strategy, particularly in patients with established MS and EBV seropositivity.

Despite these strong associations, the precise mechanisms through which EBV contributes to MS remain poorly understood. EBV may promote pathogenesis through multiple mechanisms, including molecular mimicry, dysregulated B-cell functions, amplification of autoreactive lymphocytes, and disruption of host processes via virus–host protein–protein interactions (PPIs) ^1,9–12^. Importantly, EBV’s influence may not be static but clinical type-dependent, differentially shaping neuroinflammation, demyelination, and neurodegeneration over the disease course. A critical question therefore remains: *how does EBV mechanistically contribute to the initiation and progression of MS?*

MS evolves through four clinical types. Clinically Isolated Syndrome (CIS) is defined as a first episode of inflammatory demyelination in the central nervous system (CNS), which may evolve into MS if further disease activity develops. Relapsing–Remitting MS (RRMS) presents with discrete episodes of neurological dysfunction (relapses) followed by periods of partial or full recovery (remissions), with no continuous progression between episodes. Secondary Progressive MS (SPMS) emerges after an initial RRMS course and is characterized by gradual, sustained neurological decline, sometimes accompanied by additional relapses. In contrast, Primary Progressive MS (PPMS) is marked by steady neurological deterioration from disease onset, without an initial relapsing–remitting phase ^13^. Clinically, MS often follows a sequence in which CIS progresses to RRMS and, over time, transitions into SPMS, whereas PPMS represents a distinct disease course characterized by gradual progression from onset without preceding relapses ^13^.

The RRMS clinical type is particularly important, as its cyclical pattern mirrors the life cycle of latent herpesviruses such as EBV, which persist in host cells by evading immune surveillance but can periodically reactivate. Environmental triggers, including stress and temperature fluctuations, are known to induce viral reactivation ^14–17^. Consistently, relapses in RRMS occur two to three times more frequently during or shortly after viral infections ^18–24^. These observations suggest that viral exposure contributes to increased disease activity in RRMS, supporting a role for viruses in modulating the inflammatory phase of the disease. By contrast, this viral association appears to diminish in the progressive clinical types (SPMS and PPMS), where pathology and clinical features diverge from the relapsing–remitting course ^25–27^. RRMS is primarily driven by inflammation-mediated demyelination, leading to axonal damage and lesion formation ^26^. Over time, repeated inflammatory episodes accumulate, ultimately driving progression to SPMS, which is characterized by sustained and irreversible neurological decline ^26^. Unlike RRMS, the progressive types are marked by the absence of new inflammatory lesions and limited responsiveness to immunomodulatory therapies ^26,27^. Together, these observations support a model in which viral infections, particularly EBV, exert their greatest influence during the early inflammatory phase of MS (CIS and RRMS), while their impact appears reduced in the later neurodegenerative types. Clarifying EBV’s clinical type– specific role is therefore critical for guiding the development of virus-targeted interventions tailored to distinct phases of the disease.

In earlier work, Onisiforou *et al*. (2021) introduced an integrative network-based bioinformatics framework that combined virus–host PPIs with disease-associated proteins to uncover viral-mediated mechanisms contributing to the onset and progression of neurodegenerative diseases. While initially applied to MS and viruses implicated in its development, including EBV ^1^, the framework was later extended to other contexts such as COVID-19, additional neurodegenerative disorders (e.g., Parkinson’s Disease (PD) and Alzheimer’s Disease (AD)), neuropsychiatric disorders, and diverse viral species, yielding system-level insights into how pathogens perturb host molecular networks ^28,29^. Extensions to microbiota–immune interactions and microbiota–virus crosstalk further highlighted the complexity of immune dysregulation in neurodegeneration ^30^. Although these studies provided important insights, the approach was constrained by its reliance on static protein interaction data, lacking integration with transcriptomic information and therefore unable to capture clinical type–specific dynamics. Incorporating transcriptomic analyses helps overcome this limitation by revealing tissue-, sex-, and type-specific molecular alterations ^31–33^, thereby providing deeper insights into disease mechanisms.

Experimental approaches, while essential, also face limitations in dissecting EBV dynamics in MS. Longitudinal serological studies have firmly established EBV as a causal factor but offer little mechanistic resolution. Laboratory studies have revealed several key processes, including molecular mimicry through clonally expanded B cells cross-reacting with both EBNA1 and the glial adhesion molecule GlialCAM ^9^, as well as enrichment of EBV interactors in MS genetic risk loci ^7^. More recent immunophenotyping has shown that EBV reactivation in treatment-naïve MS patients correlates with reduced circulating B-cell proportions and expansion of cytotoxic CD56^dim^ NK cells, while cerebrospinal fluid B-cell levels remain unaffected in the absence of NK surveillance, underscoring the compartmental impact of viral reactivation ^34^. Yet, these experimental findings remain fragmented, temporally constrained, and unable to capture transcriptomic reprogramming across clinical types.

To address these challenges, we developed *VirTrack*, a novel computational framework that infers mechanisms by which EBV contributes to MS pathogenesis through modulation of virus–host interactions. Unlike prior network-based approaches, VirTrack integrates experimentally validated EBV–host PPIs with clinical type–specific transcriptomic data, enabling resolution of how viral influences shift across CIS, RRMS, SPMS, and PPMS. This allows a dynamic and mechanistic view of EBV-driven host perturbations. Crucially, such modeling of evolving, clinical type–dependent viral interactions is not attainable with current experimental methods, which cannot systematically reconstruct viral–host dynamics in vivo. *VirTrack* therefore delivers a clinical type–resolved, mechanistically grounded, and experimentally inaccessible perspective on EBV-mediated pathogenesis. More broadly, it provides a generalizable framework for uncovering viral influences across viral-associated diseases, with direct implications for clarifying etiology and identifying therapeutic targets that would otherwise remain hidden.

## 2. Results

### 2.1 Transcriptomic Signatures across MS Clinical Types

Differential expression analysis of peripheral blood transcriptomes across the four clinical types of MS revealed marked transcriptional alterations, with the largest number of differentially expressed genes (DEGs) observed in the inflammatory types. CIS exhibited 2,263 DEGs (1,052 upregulated and 1,211 downregulated), while RRMS showed a comparable profile with 2,118 DEGs (995 upregulated and 1,123 downregulated). By contrast, the progressive types displayed substantially fewer DEGs, with 347 in SPMS (204 upregulated and 143 downregulated) and 615 in PPMS (327 upregulated and 288 downregulated).

### 2.2 *VirTrack* Reveals Clinical Type-Specific EBV-Driven Molecular Influences through the Integration of EBV–Host PPIs with Clinical Type-Specific MS Transcriptomic Data

*VirTrack* enables clinical type-specific mechanistic inference of EBV-driven host perturbations by integrating experimentally validated EBV–host PPIs with MS clinical type-specific transcriptomic data. For each clinical type, MS clinical type-specific DEGs were combined with high-confidence STRING host–host PPIs and EBV–host PPIs to construct integrated EBV–host–MS clinical type-specific PPI networks. These type-specific networks represent the molecular interfaces, both direct and indirect, through which EBV may perturb disease processes in each clinical type. This dual-layer integration allows, for the first time, inference of which MS pathways are perturbed by EBV at the transcriptomic level, and how these perturbations differ between early inflammatory and progressive clinical types.

#### 2.2.1 Clinical Type-Specific Interactions of EBV on Upregulated and Downregulated DEGs in MS

Using the EBV–host–MS clinical type–specific PPI networks, we extracted the EBV–host interactions associated with upregulated and downregulated DEGs in each clinical type to identify relevant viral targets and assess how these interactions change across MS. To account for differences in transcriptome size between clinical types, the number of EBV-interacting genes was normalized to the total number of DEGs per clinical type. This analysis revealed a distinct interaction pattern across MS clinical types. In CIS, EBV targeted 308 DEGs in total, corresponding to 13.6% of all DEGs. This included 207 upregulated DEGs (19.7% of all upregulated genes) and 101 downregulated DEGs (8.3% of all downregulated genes), indicating a strong upward bias (**Figure 2**). In RRMS, EBV interacted with the largest number of host genes (378 DEGs; 17.8% of all DEGs), comprising 190 upregulated DEGs (19.1%) and 188 downregulated DEGs (16.7%), reflecting a more balanced interaction profile. In SPMS, EBV connectivity was lowest, engaging only 36 DEGs (10.4% of all DEGs), including 20 upregulated (9.8%) and 16 downregulated (11.2%) genes, consistent with a diminished viral–host signature in the progressive phase. In PPMS, EBV targeted 103 DEGs (16.7% of all DEGs), consisting of 64 upregulated DEGs (19.6%) and 39 downregulated DEGs (13.5%), suggesting a narrower but still upward-biased viral influence. Collectively, these findings demonstrate that EBV–host interactions are clinical type–dependent: CIS shows a pronounced upward skew, RRMS reaches the peak of connectivity with balanced interactions, SPMS displays minimal engagement, and PPMS retains a selective upward bias.

**Figure 1:**
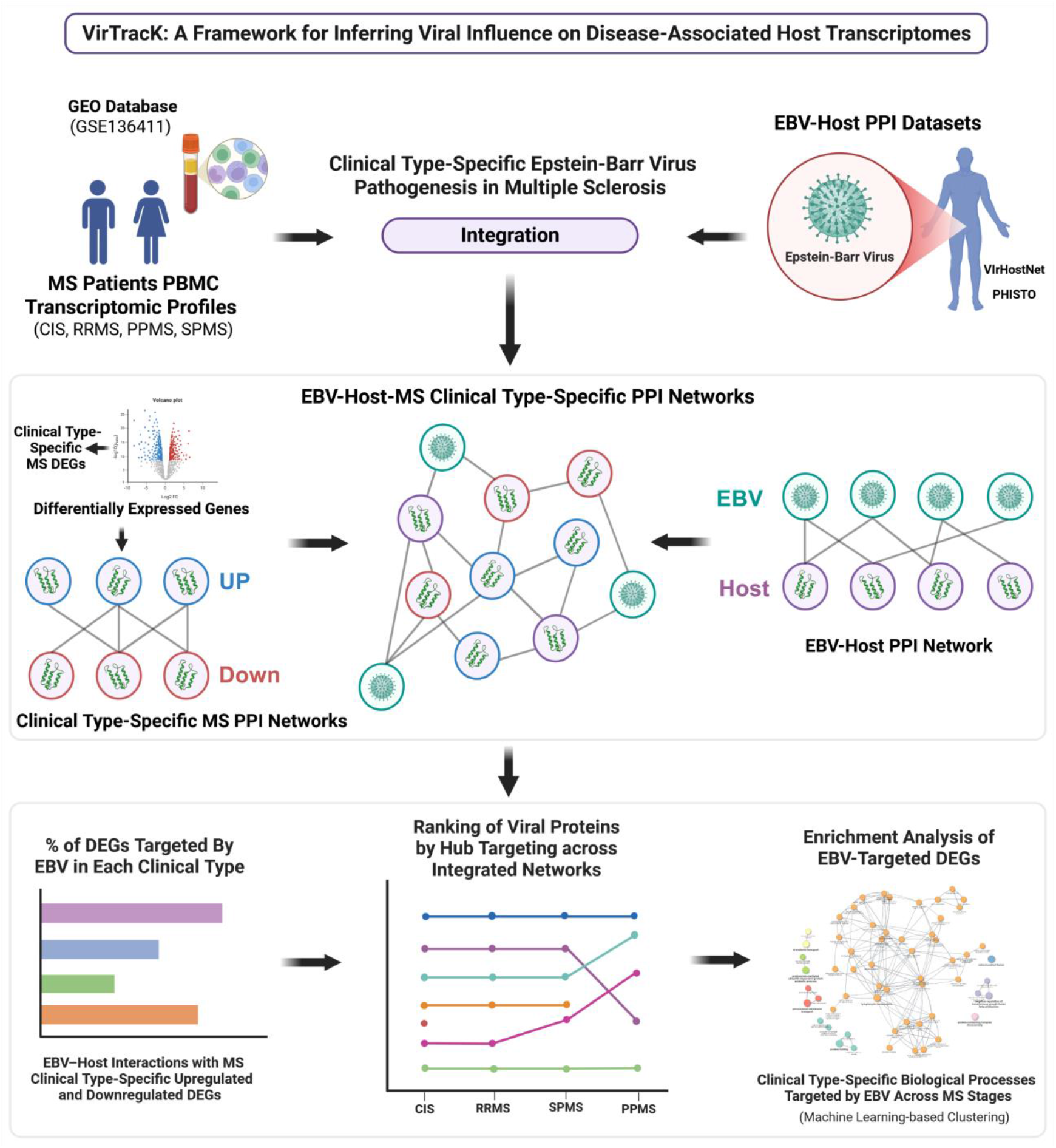
Schematic representation of the VirTrack framework applied to infer clinical type–specific EBV influences in MS. Transcriptomic profiles from MS patients (CIS, RRMS, SPMS, PPMS) obtained from the GEO database were analyzed to identify clinical type-specific differentially expressed genes (DEGs), which were then integrated with the EBV–host PPI network to construct EBV–host–MS clinical type-specific PPI networks. Subsequent analyses identified: (i) the percentage of DEGs targeted by EBV in each clinical type and specific viral protein interactions with upregulated and downregulated DEGs across MS clinical types, (ii) the ranking of viral proteins by hub-targeting activity within each EBV–host–MS clinical type-specific PPI network, and (iii) the enrichment of biological processes selectively targeted by EBV across MS clinical types, including functional clustering using machine learning approaches. Figure created with Biorender.

**Figure 2.**
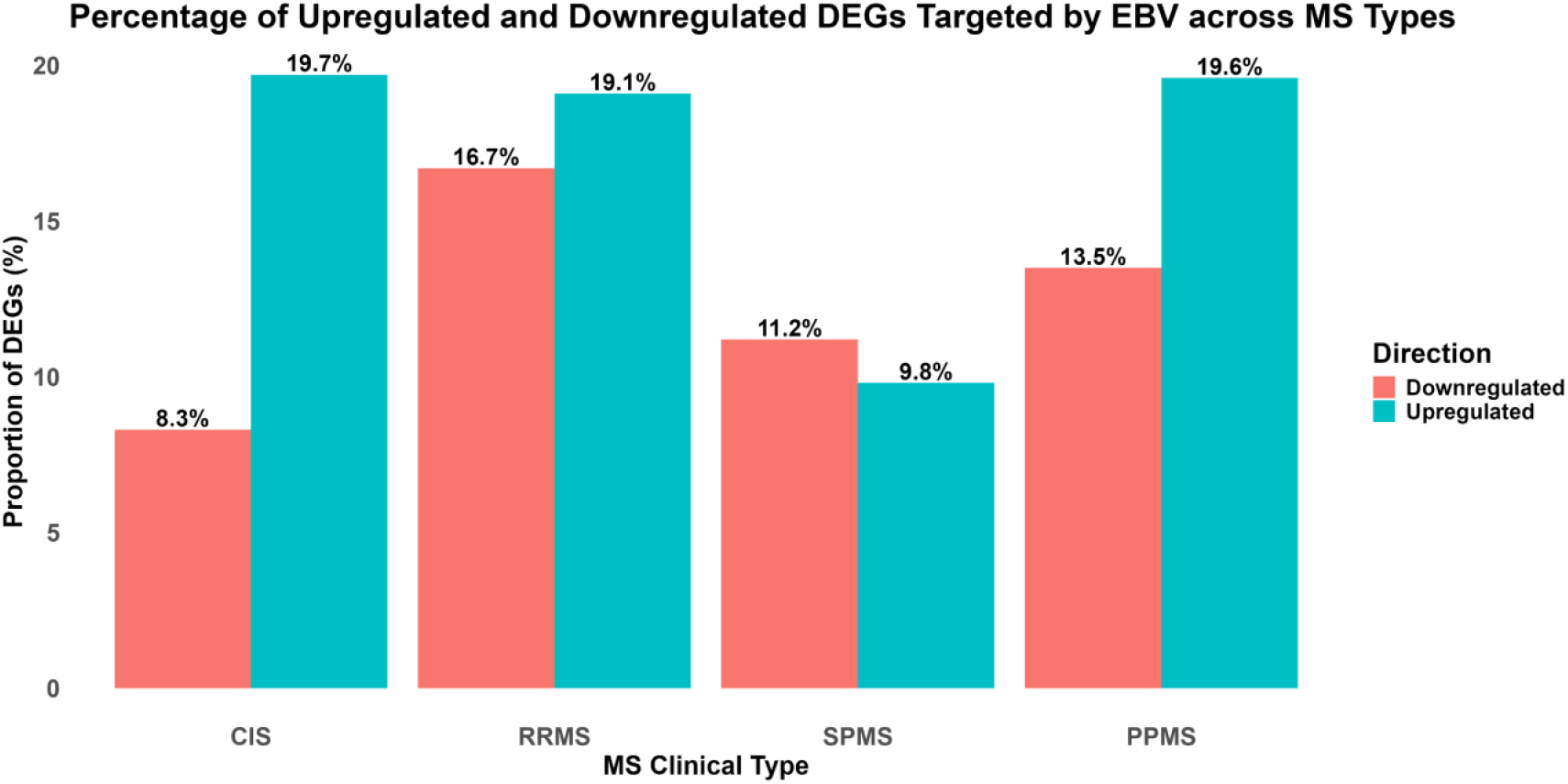
Percentage of host DEGs targeted by EBV across MS clinical types. Bar plots show the proportion of upregulated (orange) and downregulated (blue) DEGs that interact with EBV proteins in CIS, RRMS, SPMS, and PPMS. Values above the bars indicate the percentage of clinical type-specific DEGs engaged by EBV.

Further analysis of clinical type-specific EBV–host interactions highlighted several viral proteins that consistently dominated across clinical types (**Figure 3**). Our clinical type-specific analysis identified a set of EBV proteins that consistently interacted with dysregulated DEGs across MS clinical types. In CIS, the most connected proteins were BNLF2A and BVLF1, each linked to 29 DEGs, with additional contributions from BDLF4 and BHRF1. In RRMS, BNLF2A showed the broadest range of interactions, associated with 41 DEGs (23 upregulated and 18 downregulated), while BVLF1 and BDLF4 also remained prominent with 34 targeted DEGs. In SPMS, EBV–host connectivity overall was limited, with no single protein showing strong dominance, with BHRF1, EBNA-LP and BNLF2A being in the top 3. In PPMS, both BNLF2A and BVLF1 continued to appear as the top proteins, although with a reduced number of interacting DEGs (12 each), reflecting the narrower transcriptomic activity at this clinical type.

**Figure 3:**
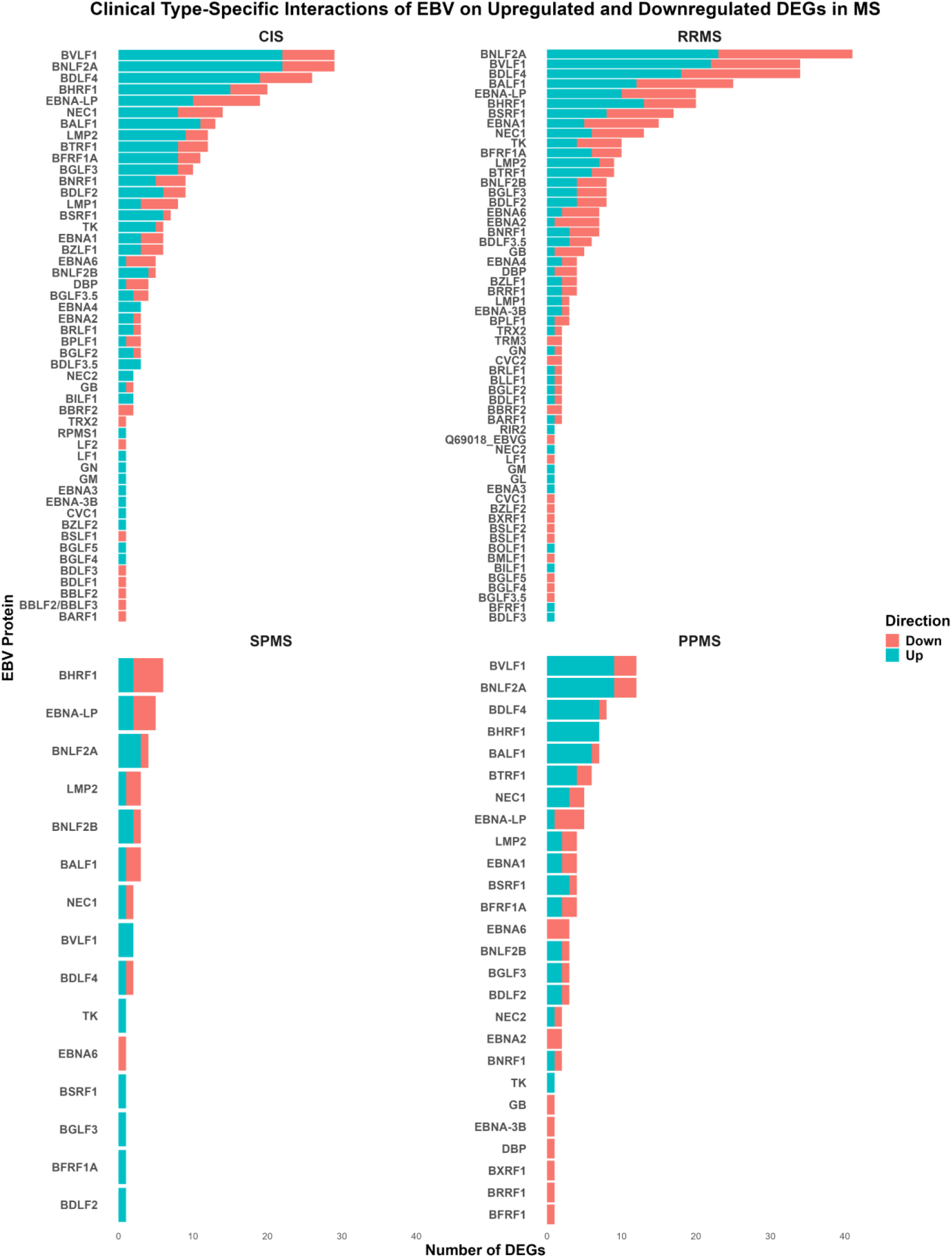
Clinical type-specific interactions of EBV with upregulated and downregulated DEGs in MS. Bar plots illustrate the number of EBV–host PPIs that interact with upregulated (orange) and downregulated (blue) DEGs in CIS, RRMS, SPMS, and PPMS.

Comparison across clinical types highlights BNLF2A and BVLF1 as the most persistent interactors, appearing among the top proteins in CIS, RRMS, and PPMS. These proteins are known to function in immune evasion (BNLF2A) and transcriptional regulation (BVLF1), which may explain their consistent association with dysregulated host genes. LMP1 was more pronounced in CIS and RRMS but less evident in progressive clinical types, suggesting that they may play a greater role in early and relapsing phases of disease. Importantly, these results do not indicate whether EBV activity is responsible for the observed up- or down-regulation, but rather show that EBV proteins directly target genes that are dysregulated in a clinical type-specific manner.

#### 2.2.2 EBV Proteins Exhibit Clinical Type-Specific Targeting of Host Hubs in the Integrated EBV–Host–MS PPI Networks

To assess whether EBV proteins preferentially target highly connected host nodes, we ranked all viral proteins by their engagement with hubs in each of the integrated EBV–host–MS clinical type-specific PPI networks. This approach enabled us to identify viral proteins exerting disproportionate influence on the structural backbone of host networks and to track whether these patterns were stable or dynamic across MS clinical types. By combining the total number of human targets with the subset corresponding to hubs, we distinguished proteins that interact with many host proteins from those that concentrate their interactions on highly connected hubs. The composite hub-targeting scores revealed consistently dominant viral proteins as well as clinical type-dependent or weaker contributors, underscoring complementary strategies by which EBV perturbs host interactomes across MS clinical types.

A comparative slopegraph (**Figure 4A**) illustrates the relative hub-targeting ranks of EBV proteins across MS clinical types. EBNA-LP consistently emerged as the top-ranked hub engager, maintaining dominance across all four types (CIS, RRMS, SPMS, and PPMS). Several other proteins also showed stable, high hub-targeting ranks: BZLF1, BVLF1, LMP2, and BDLF4 all remained within the top five, underscoring their persistent influence on central host hubs. EBNA1, BNLF2A, BNLF2B, and BHRF1 also retained relatively high ranks with only modest fluctuations between types. In contrast, some proteins exhibited weaker or more variable hub engagement. For example, BSLF1 ranked within the top 10 in CIS but fell to rank 41 in SPMS, suggesting a restricted, clinical type-specific role. BALF1 showed moderate but steady hub targeting across MS clinical types (ranks 10–12), while proteins such as BTRF1, BSRF1, BRLF1, and BGLF3 occupied a mid-range tier with minor shifts across types. Notably, LMP1, despite its established role as an EBV oncoprotein, consistently ranked very low (40–41 in CIS and RRMS, rising only to 19 in SPMS). Rare proteins such as DUT appeared only in CIS (rank 12), consistent with transient, type-specific targeting. Together, these hub-rank patterns highlight a core set of proteins (EBNA-LP, BZLF1, BVLF1, LMP2, BDLF4) that dominate hub engagement across clinical types, contrasted by proteins such as BSLF1, DUT, and LMP1, which show limited or variable hub involvement. This dual pattern suggests that EBV sustains disease-relevant perturbations through a combination of stable hub-targeting proteins and more flexible, type-specific strategies.

**Figure 4:**
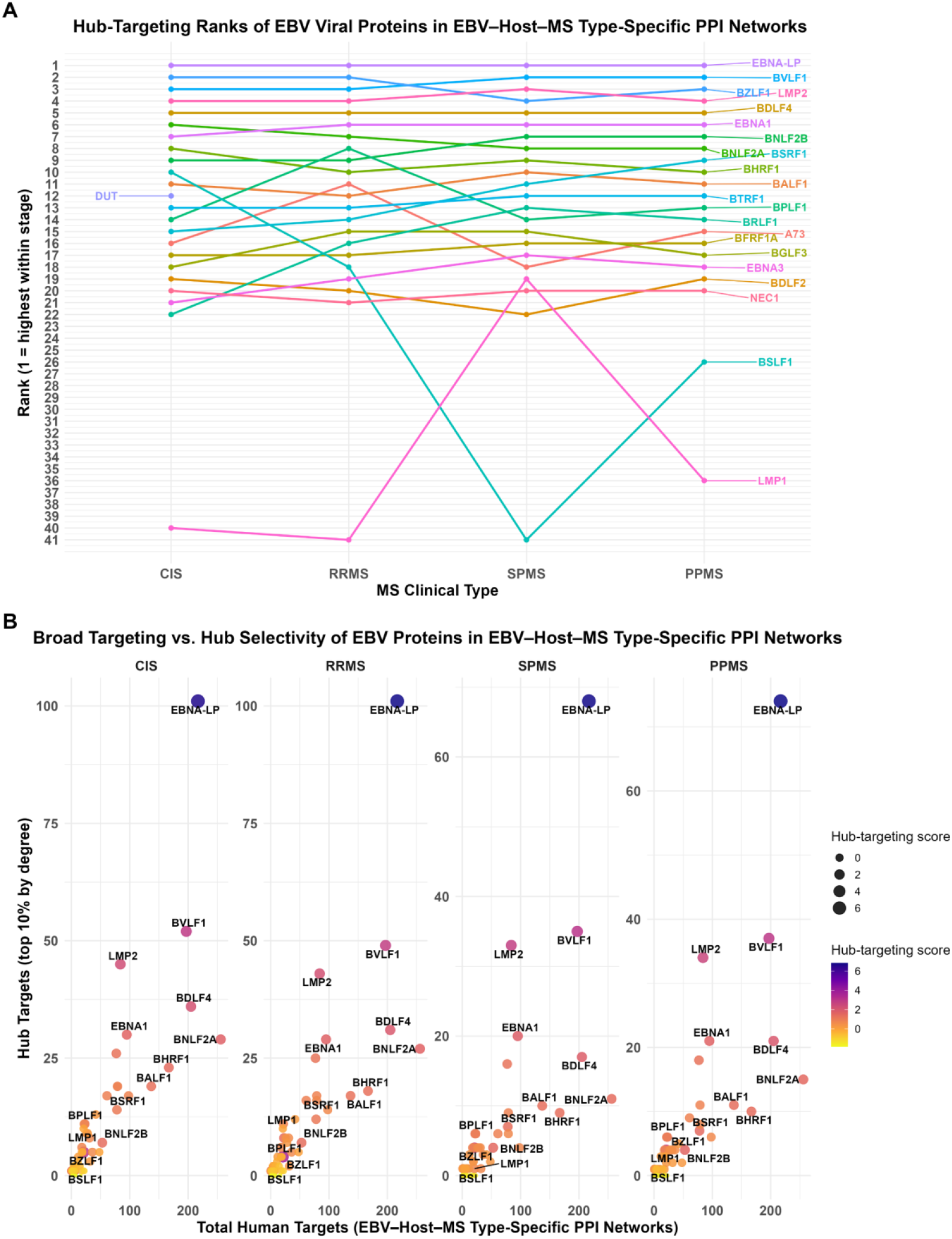
Clinical type-specific hub targeting of EBV proteins in integrated EBV–host–MS clinical type PPI networks. **(A)** Slopegraph showing the ranks of EBV proteins by hub targeting across CIS, RRMS, SPMS, and PPMS clinical types. **(B)** Scatterplots comparing total human targets (x-axis) and hub targets (y-axis) for EBV proteins across CIS, RRMS, SPMS, and PPMS clinical types, with point size and color representing hub-targeting scores across clinical types.

We then examined the balance between broad targeting and hub selectivity by comparing the total number of human proteins engaged by each EBV protein with the subset corresponding to hubs in the integrated EBV–host–MS clinical type-specific PPI networks (**Figure 4B**). This distinction allowed us to separate viral proteins that interact broadly with many host proteins from those that disproportionately concentrate their interactions on hubs. EBNA-LP was the most striking example, consistently engaging more than 200 human proteins per clinical type, including 68–101 hubs, and achieving the highest hub-targeting scores across all clinical types. BNLF2A also showed extensive targeting (~250 human proteins) but engaged fewer hubs (11-29), with its hub involvement declining in progressive clinical types, suggesting a more diffuse but less hub-focused profile. BVLF1, while narrower in scope (197 human proteins), maintained high ranks due to stable engagement with 35–52 hubs. By contrast, LMP2 interacted with relatively few total proteins (84) but disproportionately targeted network hubs, consistently connecting to one-third to half of its host targets that were hubs, thereby maintaining a top-five ranking despite its smaller target set.

Intermediate profiles were observed for proteins such as EBNA1 and BDLF4, which showed moderate effect (95 and 205 proteins, respectively) and consistent engagement with 17–36 hubs. Other proteins, including BHRF1, BNLF2B, and BALF1, displayed weaker and more variable hub targeting: although they engaged moderate numbers of proteins (50–170), only 7–23 were hubs, yielding lower overall scores. Still, they ranked within the top 10 in certain clinical types, highlighting context-dependent roles. In contrast, LMP1, despite being a well-characterized EBV oncogene, consistently ranked low due to targeting very few host proteins. Together, these findings reveal that EBV proteins employ two complementary modes of network perturbation: (i) extensive targeting, covering both hubs and non-hubs (e.g., EBNA-LP, BNLF2A, BVLF1), and (ii) hub-focused targeting, where fewer total proteins are engaged but they are disproportionately central (e.g., BZLF1, LMP2). This division of labor underscores how EBV balances robustness with specificity to sustain host network perturbations across MS clinical types.

#### 2.2.3 Biological Processes Targeted by EBV Across MS Clinical Types

We next performed Gene Ontology (GO) Biological Processes (BP) enrichment analysis on the sets DEGs that are directly targeted by EBV viral proteins in each MS clinical type, analyzing upregulated and downregulated genes separately to identify the functional pathways most impacted by viral–host interactions. To reduce redundancy and reveal higher-order patterns, enriched terms were further organized using spectral clustering, which consolidated related processes into coherent functional groups for each clinical type.

##### 2.2.3.1 EBV-Targeted Upregulated Processes in CIS: Remodeling of Immune Homeostasis, B Cell Biology, and Metabolic Programs

Enrichment analysis of EBV-targeted upregulated DEGs in the CIS clinical type identified 59 significantly enriched GO BP terms. Clustering of the upregulated terms yielded eight functional groups (**Figure 5A/B**). The dominant cluster, accounting for 71.2% of all terms, was linked to *lymphocyte homeostasis* and encompassed processes such as *B cell activation, B cell homeostasis, regulation of Toll-like receptor signaling, cellular response to molecules of bacterial origin*, and *response to fungus*. This enrichment suggests that EBV may exert early effects on both adaptive and innate immunity, particularly through modulation of B cell biology and pathogen-sensing pathways.

**Figure 5:**
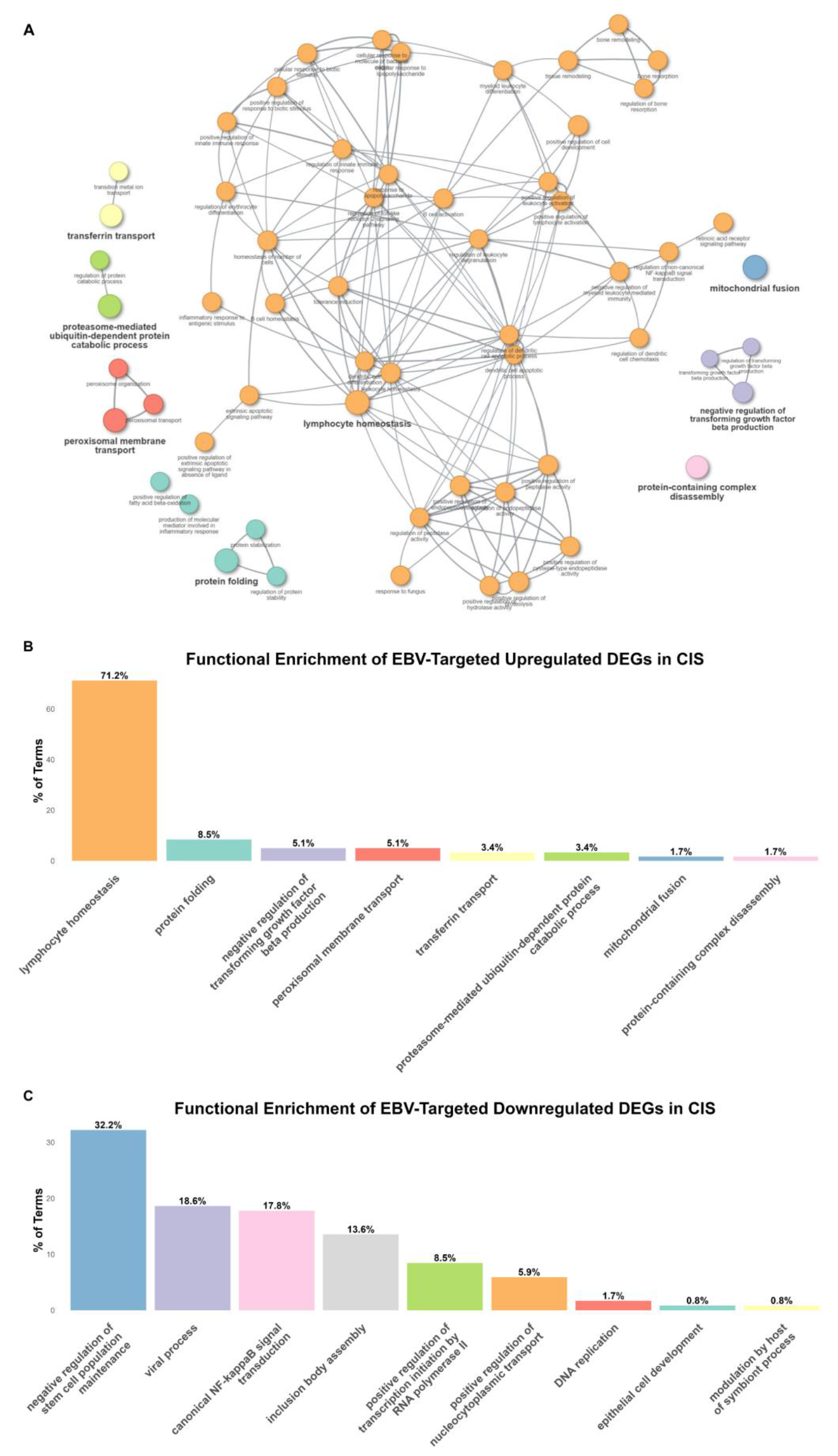
Functional enrichment of EBV-targeted DEGs in the CIS clinical type. **(A)** Spectral clustering network of significantly enriched GO BP among EBV-targeted upregulated DEGs in CIS, with nodes representing terms, clustered into groups, and edges weighted by κ-score similarity. **(B)** Bar plot of functional groups enriched among EBV-targeted upregulated DEGs in CIS, showing the percentage of total enriched terms per group. **(C)** Bar plot of functional groups enriched among EBV-targeted downregulated DEGs in CIS, showing the percentage of total enriched terms per group.

Smaller clusters reflected additional immune and metabolic processes. *Protein folding* (8.5%) and *proteasome-mediated ubiquitin-dependent catabolism* (3.4%) highlighted viral manipulation of protein quality control and host proteostasis. The cluster related to *negative regulation of TGF-β production* (5.1%) pointed toward immunometabolic reprogramming under viral influence. Organelle-related processes included *peroxisomal membrane transport* (5.1%) and *mitochondrial fusion* (1.7%), suggesting EBV engagement with organelle dynamics and cellular stress responses. Additional clusters included *transferrin transport* (3.4%) and *protein-containing complex disassembly* (1.7%), processes that may intersect with viral strategies for iron handling and disruption of host protein machinery. Together, these findings indicate that in CIS, EBV primarily interfaces with immune homeostasis, particularly B cell and innate receptor pathways, while also perturbing essential metabolic, organelle, and proteostatic programs. This suggests that EBV simultaneously shapes host immunity and cellular stress responses during the earliest clinical type of MS.

##### 2.2.3.2 EBV-Targeted Downregulated Processes in CIS: Impaired Stem Cell Maintenance, NF-κB Signaling, and Viral Response Processes

Enrichment analysis of EBV-targeted downregulated DEGs in the CIS clinical type identified 118 significantly enriched GO BP terms. Clustering of these terms yielded 10 functional groups (**Figure 5C**). The largest cluster, comprising 32.2% of terms, was linked to *negative regulation of stem cell population maintenance*, and included processes such as *response to IL-17, cellular response to IL-17, positive regulation of astrocyte differentiation*, and *positive regulation of gliogenesis*, suggesting that EBV targeting in CIS may impair regenerative and repair programs in the CNS. A second major cluster, *viral process* (18.6%), encompassed host pathways broadly involved in virus–host interactions, consistent with EBV’s capacity to co-opt cellular machinery for persistence.

The *canonical NF-κB signaling* cluster (17.8%) included both *MyD88-dependent* and *MyD88-independent Toll-like receptor (TLR) signaling processes*, highlighting EBV’s ability to disrupt innate immune sensing and transcriptional control of inflammation at multiple levels. Another major cluster, *inclusion body assembly* (13.6%), incorporated several terms related to *ubiquitin-dependent protein catabolic processes*, including *regulation of ubiquitin-dependent catabolism* and *proteasome-mediated ubiquitin-dependent catabolic processes*. This suggests that EBV interference with host proteostasis may extend beyond protein aggregation to direct manipulation of the ubiquitin–proteasome system, a pathway central to both antiviral defense and neuronal homeostasis. Other enriched groups included *positive regulation of transcription initiation by RNA polymerase II* (8.5%) and *positive regulation of nucleocytoplasmic transport* (5.9%), pointing to viral engagement with nuclear trafficking and gene expression regulation. Smaller clusters captured diverse processes, such as *DNA replication, epithelial cell development*, and *modulation by host of symbiont processes*. Collectively, these findings suggest that in CIS, EBV targeting of downregulated DEGs compromises regenerative signaling and transcriptional control, particularly through suppression of stem cell and NF-κB/TLR pathways, while simultaneously engaging host proteostasis and persistence mechanisms.

##### 2.2.3.3 EBV-Targeted Upregulated Processes in RRMS: Hormone Signaling, Endocytosis, and Symbiont Entry Mechanisms

Enrichment analysis of EBV-targeted upregulated DEGs in RRMS identified 21 significant GO BP terms, which clustered into 10 functional groups (**Figure 6A**). The largest cluster (23.8%) involved cellular *responses to endogenous stimuli*, suggesting EBV engagement with broad host signaling programs. A related cluster, *hormone-mediated signaling* (4.8%), reinforced viral modulation of systemic communication networks during relapsing disease. A second major cluster, symbiont entry into host (19.0%), directly reflected viral exploitation of host cell invasion and persistence mechanisms, while another prominent cluster, *endocytosis* (14.3%), highlighted EBV’s reliance on vesicular transport and trafficking machinery.

**Figure 6:**
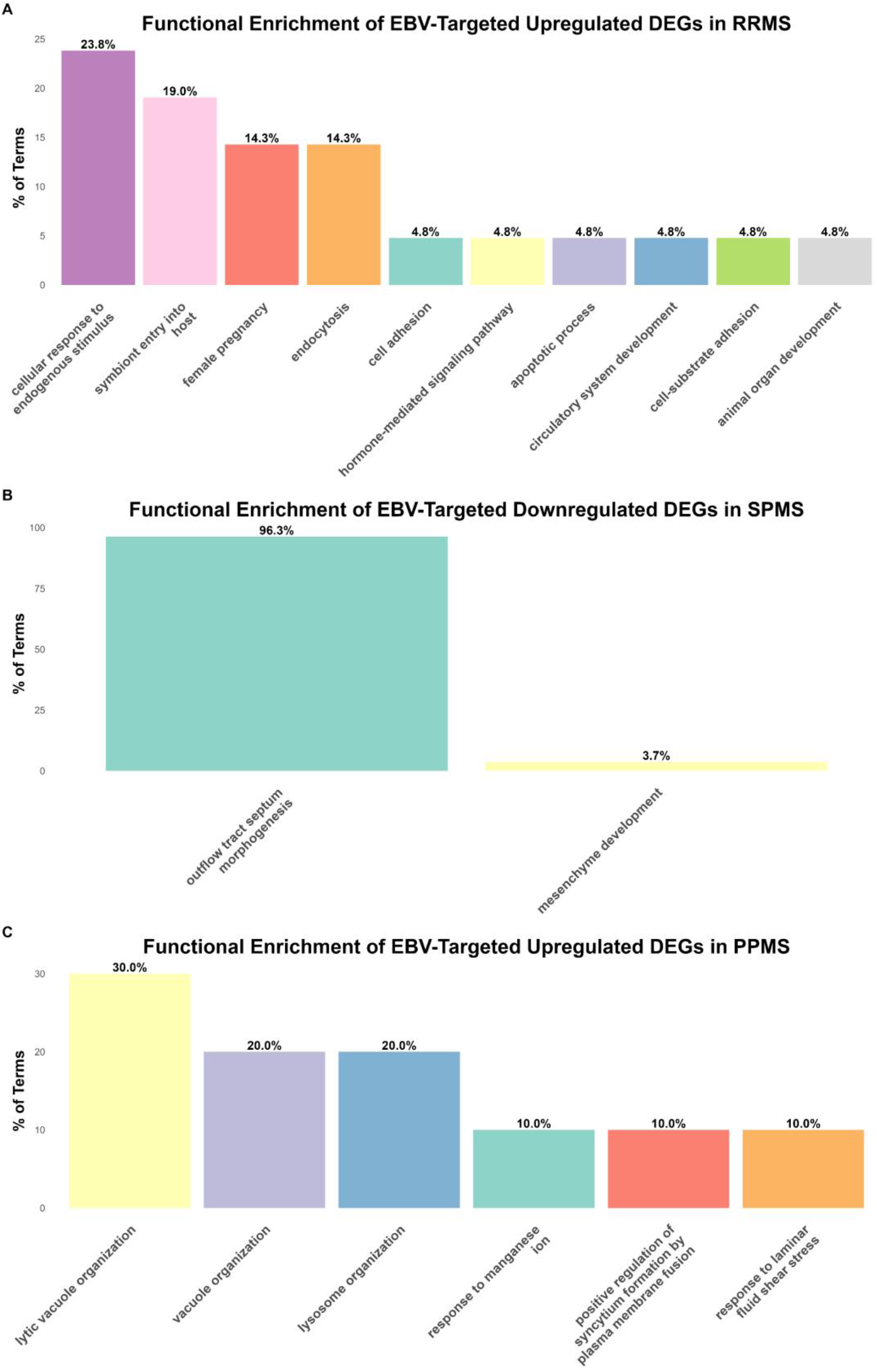
Functional enrichment of EBV-targeted DEGs in the RRMS, PPMS and SPMS clinical types. **(A)** Bar plot of functional groups enriched among EBV-targeted upregulated DEGs in RRMS, showing the percentage of total enriched terms per group **(B)** Bar plot of functional groups enriched among EBV-targeted downregulated DEGs in SPMS, showing the percentage of total enriched terms per group. **(C)** Bar plot of functional groups enriched among EBV-targeted upregulated DEGs in PPMS, showing the percentage of total enriched terms per group.

Additional smaller clusters captured diverse processes, including *female pregnancy* (14.3%), *apoptotic regulation* (4.8%), *cell adhesion* (4.8%), *cell–substrate adhesion* (4.8%), *circulatory system development* (4.8%), and *animal organ development* (4.8%). Together, these findings indicate that in RRMS, EBV-targeted upregulated pathways converge on host signaling, vesicular trafficking, and viral entry mechanisms, while additionally intersecting with adhesion, vascular, and hormone-linked processes that may influence the inflammatory lesion dynamics characteristic of relapsing disease.

##### 2.2.3.4 EBV-Targeted Downregulated Processes in RRMS: Macromolecule Localization

Enrichment analysis of EBV-targeted downregulated DEGs in RRMS yielded a single significant GO biological process term, *macromolecule localization*.

##### 2.2.3.5 EBV-Targeted Downregulated Processes in SPMS: Impaired Cardiac and Vascular Repair Pathways

Enrichment analysis of EBV-targeted downregulated DEGs in SPMS identified 27 significant GO BP terms, which clustered into two functional groups (**Figure 6B**). The dominant cluster (96.3%) was annotated to *outflow tract septum morphogenesis* and related processes, including BP terms such as *endocardial cushion morphogenesis, aortic valve morphogenesis and development, ventricular and cardiac septum morphogenesis*, and *artery/aorta development*. While these categories are classically associated with embryonic cardiogenesis, in adult biology many of these pathways are repurposed for cardiac and vascular tissue repair, endothelial–mesenchymal transition, and remodeling. Thus, their downregulation in PBMCs from SPMS patients may indicate impaired systemic programs relevant to vascular and cardiac repair. A smaller cluster (3.7%) corresponded to *mesenchyme development*, suggesting additional suppression of mesodermal differentiation pathways. No significant terms were identified for EBV-targeted upregulated DEGs in SPMS.

##### 2.2.3.6 EBV-Targeted Upregulated Processes in PPMS: Vacuolar and Lysosomal Pathways, Cell Fusion, and Stress Responses

Enrichment analysis of EBV-targeted upregulated DEGs in PPMS identified 10 significant GO BP terms, which clustered into six functional groups (**Figure 6C**). The largest clusters were related to *lytic vacuole organization* (30.0%), *vacuole organization* (20.0%), and *lysosome organization* (20.0%), pointing to a strong viral interface with host degradative and trafficking machinery. A smaller group highlighted *positive regulation of syncytium formation by plasma membrane fusion* (10.0%), consistent with EBV’s known capacity to manipulate host cell fusion processes during infection. Additional enriched pathways included *response to manganese ion* (10.0%) and *response to laminar fluid shear stress* (10.0%), suggesting possible viral interactions with metal ion homeostasis and mechanosensitive signaling. Together, these findings suggest that in PPMS, EBV preferentially engages host vacuolar and lysosomal systems while also targeting cell fusion and stress response pathways, processes that may contribute to chronic viral persistence and progressive tissue remodeling.

##### 2.2.3.7 EBV-Targeted Downregulated Processes in PPMS: Viral Gene Expression

Enrichment analysis of EBV-targeted downregulated DEGs in PPMS yielded a single significant GO biological process term, *viral gene expression*.

## 4. Discussion

Building on my earlier network-based framework ^1^, which integrated virus–host–disease PPIs with disease-associated proteins to uncover viral-mediated perturbations in neurodegenerative diseases, *VirTrack* represents a substantial methodological advance that defines a new direction for viral–host–disease network analysis. The original framework provided a broad, system-level perspective but lacked the capacity to resolve clinical type–specific dynamics or to incorporate gene expression directionality. VirTrack addresses these limitations by centering analyses on clinical type-specific MS DEGs, explicitly distinguishing between upregulated and downregulated transcripts, and thereby capturing the evolving transcriptomic landscape across CIS, RRMS, SPMS, and PPMS. This enables precise quantitative comparisons of EBV targeting across clinical types, a resolution not attainable with the original approach or with experimental methods alone. Furthermore, *VirTrack* integrates machine learning–based spectral clustering of GO enrichment results, reducing redundancy and consolidating terms into higher-order functional modules, which enhances mechanistic interpretability. Although demonstrated here in the case of EBV and MS, *VirTrack* is a generalizable framework that can be applied to any viral–disease association with available transcriptomic data, extending its scope well beyond neurodegeneration. Collectively, these innovations establish *VirTrack* as a clinical type-resolved, mechanistically focused, and broadly adaptable platform, marking a decisive departure from earlier systems-level approaches and opening new avenues for viral–host–disease research.

Analysis of the CIS clinical type revealed that EBV prominently targets pathways governing B cell activation, Toll-like receptor signaling, and immune homeostasis. Peripheral B cells latently infected with EBV display the molecular hallmarks of classical antigen-selected memory B cells ^35^, providing evidence that EBV exploits the memory compartment for persistence. Recent mechanistic work further shows that EBV-infected B cells acquire homing and migratory properties via CCL4–CCR1–FAK signaling, enabling diapedesis across endothelial barriers and disruption of vascular integrity ^36^. Together, these observations suggest that, even at the earliest clinically detectable clinical type of MS, EBV is positioned to reshape fundamental immune circuits, coupling B cell reprogramming with aberrant trafficking across the blood– brain barrier. This dual strategy not only promotes viral persistence but also establishes a foundation for chronic neuroinflammation, setting the stage for progressive network perturbations observed in later MS phases.

In RRMS, EBV-targeted genes were enriched for endocytosis, hormone signaling, and symbiont entry, supporting hypotheses that EBV reactivation contributes to MS relapses via immune mimicry and viral re-entry pathways ^37^. Clinically, relapses are known to increase following viral infections, particularly respiratory viruses ^38^, and our data suggest that EBV may potentiate such events by exploiting host entry and vesicular trafficking to amplify immune activation. Notably, EBV-targeted upregulated processes in RRMS also encompassed pathways associated with female pregnancy, aligning with evidence that gestational immune adaptations reduce relapse frequency and attenuate autoimmune activity in both MS and experimental autoimmune encephalomyelitis ^39^. In contrast, assisted reproduction technology (ART) treatments markedly increase relapse risk, driven by gonadotropin-releasing hormone–mediated cytokine shifts, VEGF induction, and enhanced immune cell trafficking across the blood–brain barrier ^40^. Together, these findings suggest that EBV reactivation intersects with hormonal and immune pathways to modulate relapse susceptibility in RRMS, positioning the virus as a key mediator within the broader immunoendocrine landscape of MS.

In progressive clinical types (PPMS, SPMS), *VirTrack* identified a reduction in overall EBV–host interactions but persistent targeting of specific processes, including lysosomal/vacuolar pathways in PPMS and vascular repair programs in SPMS. This is consistent with the broader observation that viral associations are strongest during inflammatory phases but diminish as neurodegeneration predominates. Nonetheless, sustained engagement of degradative and stress-response pathways suggests that EBV continues to shape cellular homeostasis even when overt inflammation wanes. In SPMS, enrichment analysis of EBV-targeted downregulated DEGs revealed processes linked to cardiac and vascular morphogenesis. These processes pathways in adult are used for vascular repair, endothelial–mesenchymal transition, and tissue remodeling. Their coordinated suppression in SPMS PBMCs suggests that EBV may attenuate systemic repair programs essential for vascular integrity and regeneration. This observation aligns with mechanistic work showing that EBV-infected B cells induce ICAM-1 on endothelial cells and disrupt barrier integrity during diapedesis ^36^, providing a plausible mechanistic bridge between viral perturbation and impaired vascular repair. Clinically, this may help explain why individuals with MS face elevated risks of cardiovascular disease (CVD), including myocardial infarction, stroke, and heart failure, with a 3.5-fold higher risk of all-cause mortality and a 1.5-fold increased risk of CVD-related mortality compared to the general population ^41^. Importantly, cardiovascular complications have also been directly associated with EBV infection itself, including myocarditis, coronary artery abnormalities, arrhythmias, and heart failure, which can progress to severe outcomes if untreated ^42^. Case-based evidence further underscores this link, with chronic active EBV patients developing fatal vascular complications such as giant coronary artery aneurysms, thrombosis, and aortic lesions years after infection ^43^. Taken together, these findings raise the possibility that EBV-mediated suppression of vascular repair mechanisms contributes not only to neurodegeneration in progressive MS but also to heightened cardiovascular vulnerability in later disease clinical types.

The clinical type-specific network analysis revealed that EBV engages host transcriptomes in a dynamic, type-dependent manner. In CIS, viral targeting showed a strong upward bias, with nearly one-fifth of all upregulated genes interacting with EBV proteins, suggesting that early disease is marked by viral reinforcement of pro-inflammatory and immune-activating pathways. RRMS represented the peak of connectivity, with balanced engagement of both up- and downregulated genes, consistent with the heightened immune perturbations characteristic of relapsing disease. By contrast, SPMS exhibited the lowest overall connectivity, with only ~10% of DEGs engaged, indicating a waning but not absent viral footprint in late disease. In PPMS, the number of interactions narrowed, but a selective upward bias persisted, reflecting more focused viral influence on degradative and stress-response programs. EBV proteins show clear clinical type–specific strategies for targeting host network hubs in MS.

Across all clinical types, EBV proteins displayed distinct strategies for host engagement. EBNA-LP consistently emerged as the dominant hub-targeting protein, supported by a stable core group (BZLF1, BVLF1, LMP2, BDLF4) that persistently targeted highly connected host nodes. In contrast, proteins such as BSLF1, DUT, and LMP1 exhibited variable or restricted hub engagement across clinical types, indicating more specialized or context-dependent functions. Mechanistically, EBNA-LP amplifies EBNA2-driven transcriptional programs, facilitates B cell transformation, and recruits host transcription factors to the viral genome ^44,45^. BZLF1 acts as a molecular switch controlling the latency-to-lytic transition ^46^, while LMP2A mimics B cell receptor signaling to sustain B cell survival during latency ^47^. Additionally, the prominence of BVLF1 and BDLF4 in early and relapsing clinical types suggests that transcriptional regulators beyond the canonical EBNA–LMP axis may contribute to relapse biology. Notably, BSLF1 ranked highly in CIS but declined sharply through RRMS to SPMS, suggesting an early, transient role in establishing viral–host network perturbations that wane as the disease progresses. This pattern implies that BSLF1 may contribute to the initial establishment of EBV–host interactions or immune dysregulation in early MS, but its functional relevance diminishes as the disease transitions toward chronic and progressive phases. Collectively, these findings indicate that EBV maintains its influence by strategically targeting host hubs and adapting its interactions across clinical types, providing a mechanistic basis for persistence and clinical type–specific pathology in MS.

Finally, the type-specific viral influences uncovered here have direct translational implications. Our results suggest that antiviral or EBV-targeted strategies may be most effective if deployed early (CIS/RRMS), when viral targeting is maximal. Several EBV-directed therapies are under active investigation. Several EBV-directed therapies are under active investigation, including EBV-specific cytotoxic T lymphocyte therapy ^48^, EBV vaccines ^49^, and repurposed antivirals ^50–52^. *VirTrack* can help prioritize which processes or viral proteins (e.g., EBNA-LP, BZLF1, BVLF1, LMP2, BDLF4) are most relevant to target at each clinical type.

## 5. Limitations and Future Directions

Several limitations should be acknowledged. First, this analysis was based on PBMC transcriptomes. While EBV establishes latency primarily in circulating B cells and modulates peripheral immune responses, PBMC-derived data may not fully capture compartmentalized CNS pathology. Direct CNS profiling, however, is typically limited to postmortem tissue, which does not reflect dynamic clinical type–specific processes and may be confounded by degradation artifacts. Second, the associations identified by *VirTrack* are correlative; although the framework reveals clinical type–linked interfaces between EBV proteins and dysregulated host pathways, causal relationships remain to be established. Future studies should aim to validate these findings using independent cohorts, datasets stratified by EBV serostatus, and complementary multi-omics approaches, including proteomics and single-cell transcriptomics, to refine and experimentally substantiate the inferred mechanisms.

## 6. Broader Implications

Finally, while applied here in the case of EBV and MS, *VirTrack* represents a broadly applicable framework for investigating viral influences across diverse diseases. EBV itself is implicated in a wide spectrum of conditions, including systemic lupus erythematosus, various lymphomas, and nasopharyngeal carcinoma ^53–55^. Extending *VirTrack* to these diseases may uncover both shared and disease-specific mechanisms of viral pathogenesis. Similarly, other neurodegenerative disorders, such as AD, PD, and Amyotrophic Lateral Sclerosis, have also been associated with viral or microbial triggers ^2,56,57^. By integrating viral–host interactions with disease-specific transcriptomic landscapes, *VirTrack* provides a scalable platform for disentangling viral contributions to complex diseases, offering new opportunities for identifying therapeutic targets and refining models of disease etiology.

## 7. Conclusion

This study demonstrates that EBV exerts clinical type–specific influences on the transcriptomic and network architecture of MS. Using the novel computational framework *VirTrack*, we show that EBV proteins engage distinct host hubs and pathways across CIS, RRMS, SPMS, and PPMS, with maximal viral targeting in early inflammatory clinical types and persistent but altered engagement in progressive types. Early disease is characterized by EBV-driven reprogramming of B cell and innate immune circuits, alongside convergence with bacterial- and microbiota-derived signals that may amplify inflammation and relapse risk. In contrast, progressive MS shows reduced viral–host interactions but continued disruption of lysosomal and vascular repair programs, with potential implications for both neurodegeneration and cardiovascular comorbidity. Together, these findings position EBV as a key driver of both immune dysregulation and systemic pathology in MS. *VirTrack* offers a generalizable framework for elucidating viral–host interactions across complex diseases through integration with disease-associated transcriptomic data, identifying EBV proteins and host processes as rational targets for therapeutic intervention and biomarker discovery.

## 8. Methods

### 8.1 Transcriptomic Datasets and EBV–Host PPI Data

The Gene Expression Omnibus (GEO) database was searched to identify transcriptomic datasets derived from blood samples that included large participant numbers and covered all four clinical types of MS as well as healthy controls. We selected the publicly available dataset GSE136411, which provides peripheral blood mononuclear cell (PBMC) transcriptomic profiles from 286 individuals: 67 healthy controls samples (HCs), 60 patients samples with CIS, 121 samples with RRMS, 26 samples with SPMS, and 35 samples with PPMS. This dataset represents one of the largest and most balanced PBMC transcriptome cohorts available for MS; by contrast, many previous studies included ≤50–100 samples and were limited to one or two clinical types. Importantly, GSE136411 uniquely spans all four clinical types (CIS, RRMS, SPMS, PPMS) alongside healthy controls, making it particularly well suited for clinical type–resolved analysis of viral– host interactions. Gene expression profiling was performed using two Illumina microarray platforms (HumanRef-8 v2 and HumanHT-12 v4). PBMCs provide accessible systemic immune readouts, and given EBV’s latency in circulating B cells and its modulation of peripheral immune responses, PBMC-derived transcriptomes offer meaningful insights into EBV-driven immune dysregulation, even if they do not fully capture compartmentalized CNS pathology.

EBV–host PPIs (7,073 interactions) were aggregated across three EBV strains (TaxIDs 10377, 82830, 10376), encompassing 1,153 human targets and 153 viral proteins. UniProt accessions were mapped to primary EBV gene/ORF names, with strain- or isoform-specific duplicates collapsed to yield 73 unique viral genes. Where no gene name was available, UniProt accessions were retained to avoid information loss.

### 8.2 Preprocessing and Differential Expression Analysis of MS Dataset

The GSE136411 dataset was pre-processed and analyzed in the R statistical environment. Expression data were normalized and log2 transformed prior to analysis. Differentially expressed genes (DEGs) were identified using the Limma package ^58^. Probe set identifiers were mapped to gene symbols according to the platform-specific annotation files; in cases where multiple probe sets corresponded to a single gene, the average expression value was calculated. Genes with an adjusted p-value < 0.05 were considered statistically significant and retained for downstream analyses.

### 8.3 VirTrack Framework: Integrating EBV–Host PPIs with MS Clinical Type-Specific DEGs via Construction of EBV–Host–MS Clinical Type-Specific PPI Networks

To mechanistically anchor EBV within MS clinical type-specific biology, we developed *VirTrack*, a computational pipeline that integrates disease DEGs with experimentally validated EBV–host PPIs and host–host interactomes. Both upregulated and downregulated DEGs identified from CIS, RRMS, SPMS, and PPMS (see Section 8.2) were used to construct clinical type-specific disease networks. For each clinical type, curated EBV–host PPIs were merged with the corresponding DEG network to generate EBV–host– MS clinical type-specific PPI networks. High-confidence host–host PPIs among these nodes were retrieved from the STRING database ^59^ (confidence ≥0.7). This integration produced clinical type-specific networks comprising (i) disease DEGs, (ii) host–host interactions, and (iii) EBV–host interactions. The EBV–host–MS clinical type-specific PPI networks were visualized using visNetwork package in R ^60^.

#### 8.3.1 Comparative Analysis of EBV-Targeted Upregulated and Downregulated DEGs in CIS, RRMS, SPMS, and PPMS

Using each of the constructed EBV–host–MS clinical type–specific PPI networks, we identified EBV–host interactions involving upregulated and downregulated DEGs to determine which viral proteins directly target disease-relevant genes in each clinical type. To quantify EBV’s impact, we calculated for each clinical type (CIS, RRMS, SPMS, PPMS): (i) the percentage of DEGs targeted by EBV, and (ii) the proportion of EBV-interacting DEGs within upregulated and downregulated gene sets.

#### 8.3.2 Assessment and Ranking of EBV Protein Interactions with Host Hubs Across the Integrated EBV– Host–MS Clinical Type-Specific Networks

For each EBV-host-MS clinical type-specific PPI network (CIS, RRMS, SPMS, PPMS), we assessed whether EBV proteins preferentially targeted highly connected host proteins. Networks were analyzed using *igraph* package in R ^61^, with human nodes scored by degree and betweenness centrality, and hubs defined as the top 10% of human proteins by degree within each clinical type. Viral–host interactions were extracted and standardized, and for each EBV protein we calculated the total number of human targets, the number of hub targets, the summed and average degree of targeted hubs, and the summed and average betweenness of targeted hubs. To integrate these measures, we derived a composite hub-targeting score consisting of the z-scored number of hub targets plus weighted contributions from the average hub degree and betweenness (0.5 weight each). EBV proteins were ranked within each clinical type according to this score.

#### 8.3.3 Functional Enrichment of EBV-Targeted DEGs Across MS Clinical Types and Machine Learning– Based Clustering

To functionally characterize the EBV-interacting DEGs identified in each MS clinical type, we performed GO BP enrichment analysis. For every clinical type (CIS, RRMS, SPMS, PPMS), EBV-targeted upregulated and downregulated DEGs were analyzed separately. Gene symbols were mapped to Entrez Gene identifiers using *clusterProfiler* ^62^ package in R with the org.Hs.eg.db annotation database, and only unique, valid IDs were retained. GO enrichment was carried out with *clusterProfiler* (enrichGO), restricting to the BP ontology and applying Benjamini–Hochberg correction. Significant terms were defined at adjusted *p*<0.05 and *q*<0.05. Where necessary, fold enrichment was calculated as the ratio of observed to expected gene counts.

To reduce redundancy among enriched terms and reveal higher-order functional themes, we constructed ClueGO-like term–term similarity networks ^63^. Pairwise functional similarity between GO terms was quantified using Cohen’s κ, calculated over the union of annotated genes. Based on sensitivity analysis across a range of κ thresholds, we set κ = 0.40, which provided an optimal balance between preserving biologically coherent groups and avoiding excessive network fragmentation. In the resulting networks, nodes represented GO terms and edges represented functional similarity. GO terms were then clustered into functional groups using a spectral clustering machine learning approach applied to the κ similarity matrix. The method embeds the similarity graph into a reduced eigenspace using eigenvectors of the normalized Laplacian, followed by k-means clustering. The optimal number of clusters was selected by maximizing silhouette scores. Within each group, the representative term was defined as the GO category with the smallest adjusted *p-value*; ties were broken by highest fold enrichment. Spectral clustering was chosen because it is particularly well suited for detecting communities in graphs with weighted similarity structures, handles overlapping and weakly connected clusters more effectively than modularity-based approaches (e.g., Louvain, Walktrap), and provides more stable results than density-based methods (e.g., HDBSCAN) when applied to sparse biological networks.

## 11. Data Availability Statement

All data used in this study are publicly available from the resources and references cited within the Methods section.

## 12. Code Availability Statement

The code used in this study was developed independently by the author and constitutes the author’s intellectual property. While the method is fully described in the manuscript, the original code is not publicly available but may be shared upon reasonable request, subject to intellectual property considerations.

